# Phylogenomics of expanding uncultured environmental Tenericutes provides insights into their pathogenicity and evolutionary relationship with Bacilli

**DOI:** 10.1101/2020.01.21.914887

**Authors:** Yong Wang, Jiao-Mei Huang, Ying-Li Zhou, Alexandre Almeida, Robert D. Finn, Antoine Danchin, Li-Sheng He

## Abstract

The metabolic capacity, stress response and evolution of uncultured environmental Tenericutes have remained elusive, since previous studies have been largely focused on pathogenic species. In this study, we expanded analyses on Tenericutes lineages that inhabit various environments using a collection of 840 genomes. Several novel environmental lineages were discovered inhabiting the human gut, ground water, bioreactors and hypersaline lake and spanning the Haloplasmatales and Mycoplasmatales orders. A phylogenomics analysis of Bacilli and Tenericutes genomes revealed that some uncultured Tenericutes are affiliated with novel clades in Bacilli, such as RF39, RFN20 and ML615. Erysipelotrichales and two major gut lineages, RF39 and RFN20, were found to be neighboring clades of Mycoplasmatales. We detected habitat-specific functional patterns between the pathogenic, gut and the environmental Tenericutes, where genes involved in carbohydrate storage, carbon fixation, mutation repair, environmental response and amino acid cleavage are overrepresented in the genomes of environmental lineages. We hypothesize that the two major gut lineages, namely RF39 and RFN20, are probably acetate and hydrogen producers. Furthermore, deteriorating capacity of bactoprenol synthesis for cell wall peptidoglycan precursors secretion is a potential adaptive strategy employed by these lineages in response to the gut environment. This study uncovers the characteristic functions of environmental Tenericutes and their relationships with Bacilli, which sheds new light onto the pathogenicity and evolutionary processes of Mycoplasmatales.

**IMPORTANCE:** Environmental Tenericutes bacteria were recently discovered in numerous environments. However, our current collection of Tenericutes genomes was overrepresented by those for pathogens. Our phylogenomics study displays the relationships between all the available Tenericutes, as well as those between Tenericutes and the clades in Bacilli, which casts lights into the uncertain boundary between the environmental lineages of Tenericutes and Bacilli. By comparing the genomes of the environmental and pathogenic Tenericutes, we revealed the metabolic pathways and adaptive strategies of the Tenericutes in the different environments and hosts. We also predicted the metabolism of the two major gut lineages RF39 and RFN20 of Tenericutes, indicating their potential importance in stabilization of the gut microbiome and contribution to human health.

## INTRODUCTION

The phylum Tenericutes is composed of bacteria lacking a peptidoglycan cell wall. The most well-studied clade belonging to this phylum is Mollicutes, which contains medically relevant genera, including *Mycoplasma*, *Ureaplasma* and *Acholeplasma.* All reported mollicutes are commensals or obligate parasites of humans, domestic animals, plants and insects (1). Most studies so far have focused on pathogenic strains in the Mycoplasmatales order (which encompasses the genera such as *Mycoplasma*, *Ureaplasma*, *Mesoplasma* and *Spiroplasma*), resulting in their overrepresentation in current genome databases. However, Tenericutes can also be found across a wide and diverse range of environments. Recently, free-living *Izemoplasma* and *Haloplasma* were reported in a deep-sea cold seep and brine pool, respectively (2, 3). Based on their genomic features, the cell wall-lacking *Izemoplasma* were predicted to be hydrogen producers and DNA degraders. The *Haloplasma contractile* genome encodes actin and tubulin homologues, which might be required for its specific motility in deep-sea hypersaline lake (4). These marine environmental Tenericutes exhibit metabolic versatility and adaptive flexibility. This points out the unwanted limitation that we must take into account at present when working on isolates of marine Tenericutes representatives. The paucity of marine isolates currently available has limited further mechanistic insights.

Environmental Tenericutes might be pathogens and/or mutualistic symbionts in the gut of their host species. For example, mycoplasmas and hepatoplasmas affiliated with Mycoplasmatales play a role in degrading recalcitrant carbon sources in the stomach and pancreas of isopods (5, 6). *Spiroplasma* symbionts discovered in sea cucumber guts possibly protect the host intestine from invading viruses (7). Tenericutes were also found in the intestinal tract of healthy shallow-water fish, mussels and 305 insect specimens (8–10). Recently, over 100 uncultured Tenericutes displaying high phylogenetic diversity were discovered in human gut metagenomes (11), irrespective of age and health status. It remains to be determined whether these novel lineages found in the human gut are linked to the maintenance of gut homeostasis and microbiome function. As a consequence of the host cell-associated lifestyle, the Tenericutes bacteria show extreme reduction in their genomes as well as reduced metabolic capacities, eliminating genes related to regulatory elements, biosynthesis of amino acids and intermediate metabolic compounds that must be imported from the host cytoplasm or tissue (12). Beyond genome reduction, evolution of pathogenic Mycoplasmatales species has also been accompanied by acquisition of new core metabolic and virulence factors (13, 14). Therefore, a comparison of the genetic profiles between environmental lineages and pathogens is needed to obtain insights into the adaptation of beneficial symbionts and the emergence of new diseases.

Since Tenericutes were recently reclassified into a Bacilli clade of Firmicutes (15), the discovery of environmental Tenericutes renovates the question regarding the boundary between Tenericutes and other clades of Bacilli. RF39 and RFN20 are two novel lineages of Bacilli, reported in the gut of the humans and domestic animals (16, 17). The environmental lineages of Bacilli and Tenericutes are expected to consist in close relatives but their genetic relationship has not been studied. This is important to address, as uncultured environmental Tenericutes and Bacilli may potentially emerge as pathogens. In this study, we compiled the genomes of 840 Tenericutes and determined their phylogenomic relationships with Bacilli. By analyzing the functional capacity encoded in these genomes, we deciphered the major differences in metabolic spectra and adaptive strategies between the major lineages of Tenericutes, including the two dominant gut lineages RF39 and RFN20.

## RESULTS AND DISCUSSION

### Phylogenetic tree of 16S rRNA genes and phylogenomics of Tenericutes

We retrieved all available Tenericutes genomes from the NCBI database (April, 2019). A total of 840 genomes with ≥50% completeness and ≤10% contamination by foreign DNA were selected (Supplementary file 1). From these, 685 16S rRNA genes were extracted and clustered together when displaying at >99% identity, resulting in 227 representative sequences. Approximately 70% of the non-redundant sequences were derived from the order Mycoplasmatales (highly represented by the hominis group), which was largely composed of pathogens isolated from plants, humans and animals. Together with 33 reference sequences from marine samples, a total of 260 16S rRNA genes were used to build a maximum-likelihood (ML) tree. Using *Bacillus subtilis* as an outgroup, Tenericutes 16S rRNA sequences were divided into several clades (Fig. 1A). *Acholeplasma* and *Phytoplasma* were grouped into one clade, while *Izemoplasma* and *Haloplasma* were closer to the basal group. Tenericutes species were detected across a range of environments, including mud, bioreactors, hypersaline lake sediment, and ground water. The non-human hosts of Tenericutes included marine animals, domestic animals and fungi. Sequences isolated from fungi and mycoplasma-infected animal blood samples were associated with longer branches, indicating the occurrence of a niche-specific evolution. *Hepatoplasma* identified as a novel genus in Mycoplasmatales is also exclusively present in the gut microbiome of amphipods and isopods (5, 18). *Spiroplasma* detected in a sea cucumber gut has been described as a mutualistic endosymbiont (7), rather than a pathogen. These isolates from environmental hosts were distantly related to others in the tree, indicating a high diversity of Mycoplasmatales across a wide range of hosts and their essential role in adaptation and health of marine invertebrates. Analyses of 135 16S rRNA amplicon datasets and 141 Tara Ocean metagenomes (19) from marine waters revealed the presence of mycoplasmas from the hominis group and other sequences from the basal groups of the tree in more than 21.7% of the samples. Four of the five representative 16S rRNA sequences from the hominis group were similar (95.9%-99.3%) to that of halophilic *Mycoplasma todarodis* isolated from squids collected near an Atlantic island. The finding of the Tenericutes isolated from humans and other animal hosts in the marine samples indicates that they may be spreading possibly through sewage. The relative abundance of the twelve representative 16S rRNA genes from the marine waters was extremely low (<0.1%) in the microbial communities of the oceans. However, considering the tremendous body of marine water, the oceans harbor a massive Tenericutes population composed of undetected novel lineages. We detected two major clades of human gut lineages (hereafter referred to as HG1 and HG2) that were placed between Mycoplasmatales and Acholeplasmatales (Fig. 1A). These two lineages have been revealed recently as encompassing many previously unknown species in the human gut (11). However, their contribution to human health and the core gut microbiome stability remains unclear.

**Figure 1.**
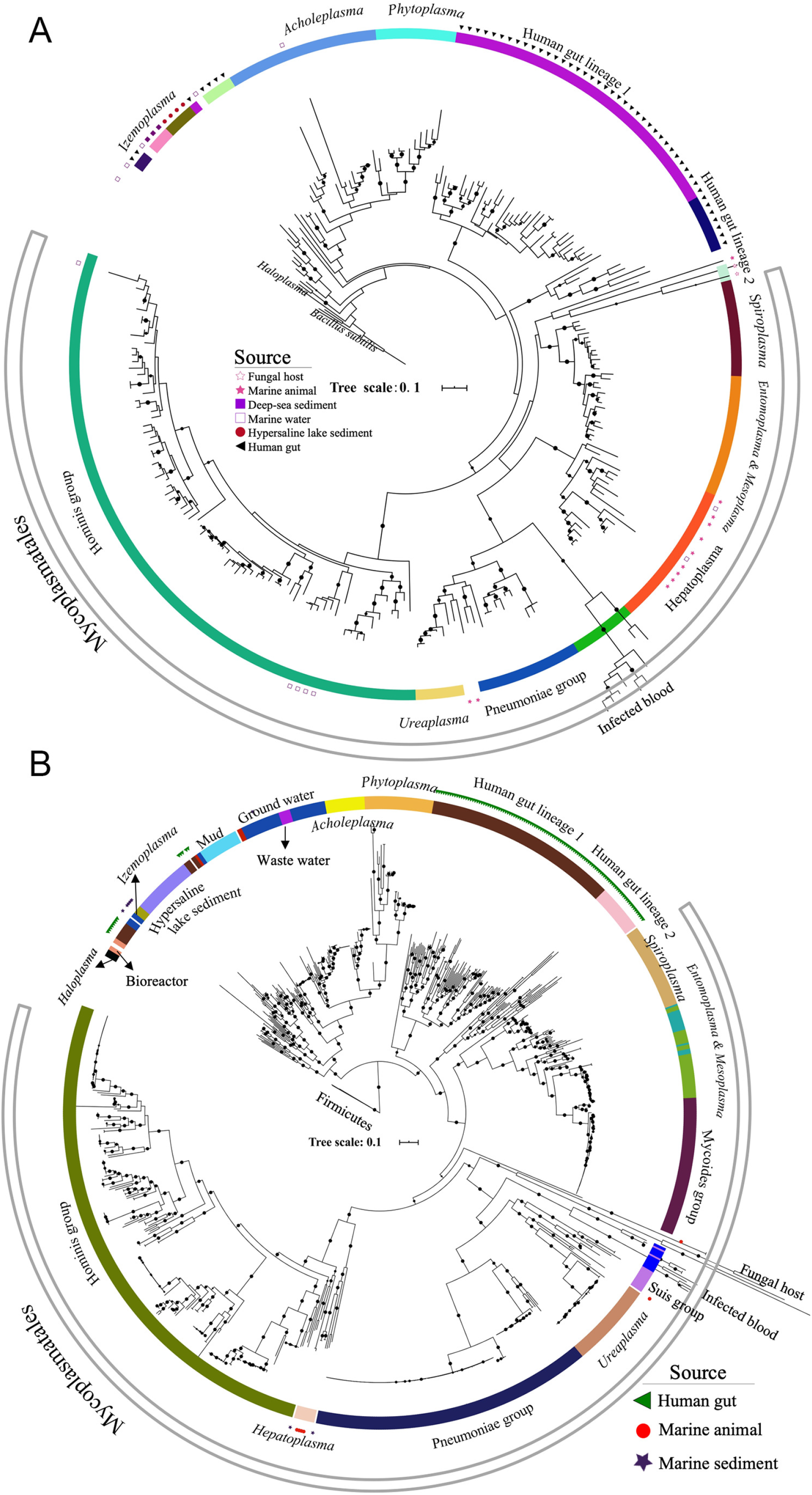
Phylogenetic trees of Tenericutes The maximum-likelihood phylogenetic trees were constructed by concatenated conserved proteins (A) and 16S rRNA genes (B). The bootstrap values (>50) are denoted by the dots on the branches.

A phylogenomics analysis of Tenericutes was performed using concatenated conserved proteins from 840 Tenericutes genomes and three Firmicutes genomes. Interestingly, the topology of the phylogenomic tree coincides with that of the phylogenetic tree based on 16S rRNA genes. However, 67.6% of the genomes were derived from Mycoplasmatales, indicating a strong bias of Tenericutes genomes towards pathogens and disease-inducing isolates. The human gut lineages HG1 (*n*=87) and HG2 (*n*=21) were found to be neighboring clades of Mycoplasmatales as well (Fig. 1B). The genetic distance between the genomes of the gut lineages was much higher than that between the species in Mycoplasmatales, except for those in mycoplasma-infected blood and fungi. *Acholeplasma* and *Phytoplasma* were within a clade composed of uncultured environmental Tenericutes lineages from ground waters, hypersaline sediments and mud, suggesting an environmental origin for the two genera.

By calculating the relative evolutionary divergence (RED) of the genomes of several Tenericutes lineages (15), the average RED values for HG1 and HG2 were 0.94±0.03 and 0.91±0.07, respectively. Considering an expected RED value of 0.92 at the genus level, these two lineages can be considered new genera in Tenericutes. The RED value for the sequences from hypersaline lake sediments was 0.70, which supports the presence of a new order or family in Tenericutes.

### Phylogenomic position of Tenericutes in Bacilli

Tenericutes were recently integrated into the Bacilli clade within the Firmicutes phylum (15). To examine the phylogenetic positions of the new Tenericutes lineages and Bacilli, we used representative genomes of the orders within Bacilli and those in Tenericutes available on NCBI. The topology of the phylogenomic relationships was supported by two ML methods. In the phylogenomic tree, four Bacilli orders, namely Staphylococcales, Exiguobacterales, Bacillales, and Lactobacillales, were clearly split from those of Tenericutes. Newly defined orders RF39, RFN20 and ML615 in Bacilli clustered with HG1, HG2, and uncultured Tenericutes from bioreactors, respectively. This suggests that most of uncultured environmental Tenericutes are probably novel Bacilli orders, and that the boundary between Tenericutes and Bacilli is uncertain. RF39, RFN20 and ML615 were also affiliated with Tenericutes if the boundary of Tenericutes on the tree was set at Haloplasmatales. Although RF39 and RFN20 are part of the HG1 and HG2 lineages, they have also been detected in domestic animals (20). Interestingly, the Erysipelotrichales order was phylogenetically placed between both human gut lineages (Fig. 2). Since all Erysipelotrichales species described in the literature so far possess a cell wall (21), their phylogenomic affinity to cell wall-lacking Tenericutes is unexpected.

**Figure 2.**
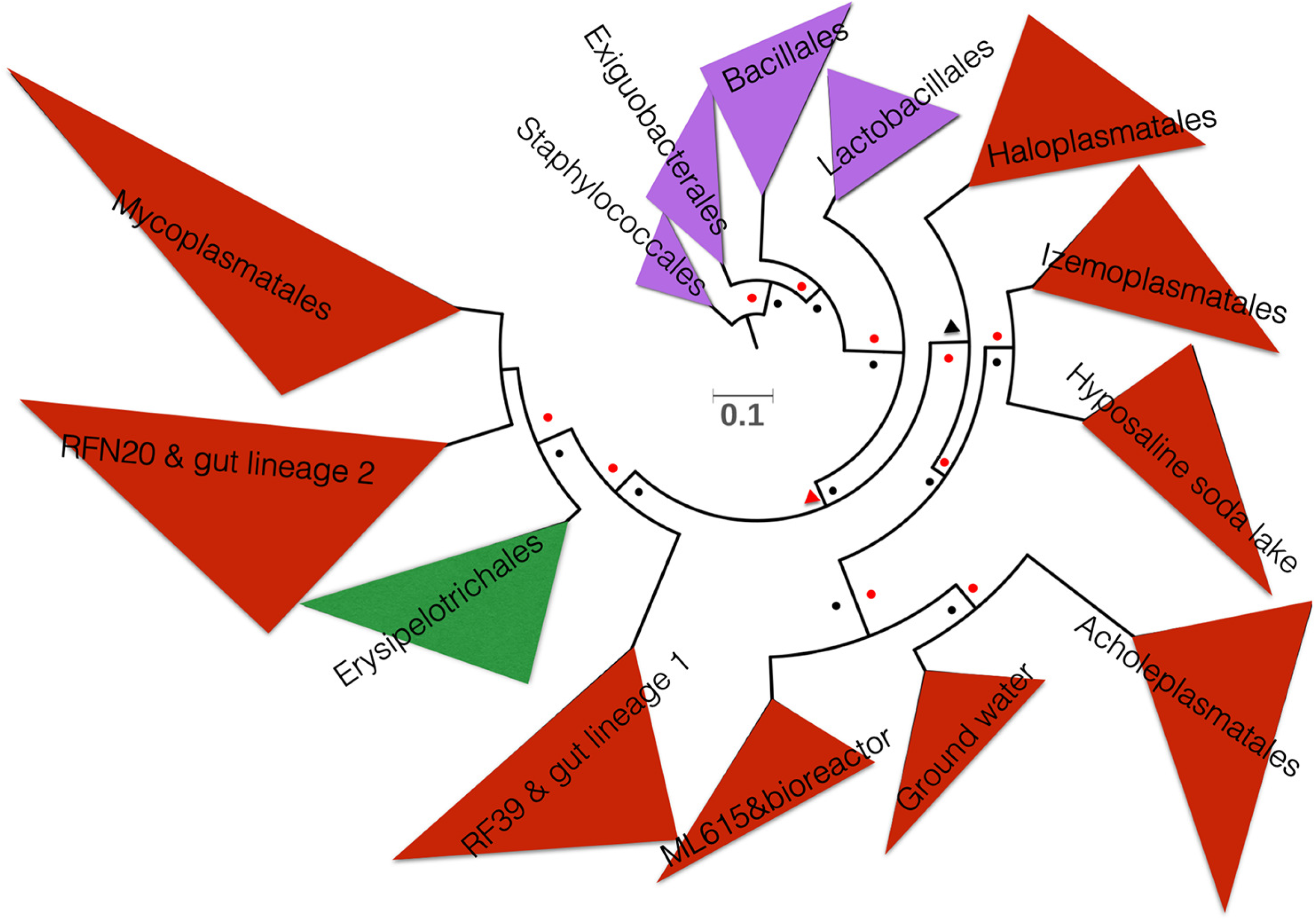
Phylogenetic positions of Tenericutes families in Bacilli. Representative genomes from orders of Bacilli were used to construct the phylogenomics tree using concatenated conserved proteins by IQ-TREE and RAxML. The bootstrap values were shown as triangles (50-90) and dots (>90) with a red color for the results of RAxML and deep blue for those of IQ-TREE, respectively. The purple clades represent the orders of Bacilli and the red ones denote Tenericutes.

We investigated the genome structure of Tenericutes and Erysipelotrichales species by calculating genome completeness, size and GC content (Fig. S1). Most of the high-quality genomes (>90% completeness and <5% contamination) were assigned to Mycoplasmatales and Acholeplasmatales. In contrast to the rather stable genomes of the pathogenic species, the genome sizes of the uncultured Tenericutes species differed from each other and almost all were smaller than 2 Mb. Haloplasmatales genomes were the largest on average. Most of the Tenericutes genomes have a low GC content (<30%), whereas the average GC content of those from a hypersaline lake was about 50%, consistent with a selection pressure exerted by ionic strength on the DNA double helix (22, 23). Notably, GC contents calculated on 1 kb intervals in Tenericutes genomes from ground water and HG1 (specifically RF39) varied from 20% to 70%, suggesting great plasticity and frequent gene transfers. However, these results were dependent on the number of genomes considered from different sources and may be influenced by the quality of genome binning.

### Genomic and functional divergence between environmental Tenericutes and pathogens

Erysipelotrichales and Tenericutes genomes were functionally annotated to characterize their metabolic pathways and stress responses that might determine the versatility and niche-specific evolution of different orders and lineages in Tenericutes. The annotation results against the Kyoto Encyclopedia of Genes and Genomes (KEGG) (24) and the clusters of orthologous groups (COGs) databases were used to calculate the percentages of the genes in the genomes (supplementary file 2). Based on the frequency of all the COGs, Erysipelotrichales and Tenericutes were split into two major agglomerative hierarchical clustering (AHC) clusters. Mycoplasmatales and *Phytoplasma* formed AHC cluster 1, while the remaining formed cluster 2.

Using Mann-Whitney test, 203 KEGG genes and 420 COGs showed a significant difference (*p*<0.01) in frequency between the two AHC clusters (supplementary file 2). We selected 62 of the genes to represent those for 16 functional categories that were distinct in environmental adaptation and carbon metabolism between the two clusters (Table S1 and Fig. 3). Sugars such as xylose, galactose and fructose might be fermented to L-lactate, formate and acetate by Tenericutes. The sugar sources and fermentation products differed between the groups (Fig. 3). Phosphotransferase (PTS) systems responsible for sugar cross-membrane transport were encoded by most of the genomes of *Spiroplasma*, *Mesoplasma*, *Entomoplasma*, Haloplasmatales, Erysipelotrichales, mycoides, and pneumoniae groups. Although most of the environmental Tenericutes genomes did not maintain PTS systems, sugar uptake might be carried out by ABC transporters. Almost all of the Tenericutes groups in the AHC cluster 2 (containing all the environmental lineages) were found to encode genes involved in starch synthesis (*glgABP*) and carbon storage, except for HG1. These Tenericutes groups also encoded the pullulanase gene PulA involved in starch degradation. Autotrophic pathways were present almost exclusively in environmental Tenericutes genomes. CO_2_ is fixed by two autotrophic steps mediated by the citrate lyase genes that function in reductive citric acid cycle (rTCA) and the 2-oxoglutarate/2-oxoacid ferredoxin oxidoreductase genes (*korABCD*) that encode enzymes for reductive acetyl-CoA pathway. The resulting pyruvate might be further stored as glucose and glycan via reversible Embden–Meyerhof–Parnas (EMP) pathway. PPDK is the key enzyme that controls the interconversion of phosphoenolpyruvate and pyruvate in prokaryotes (25). Among all the environmental lineages and Erysipelotrichales, *ppdK* gene was frequently identified (73.8%-100%) except for Haloplasmatales and Acholeplasmatales.

**Figure 3.**
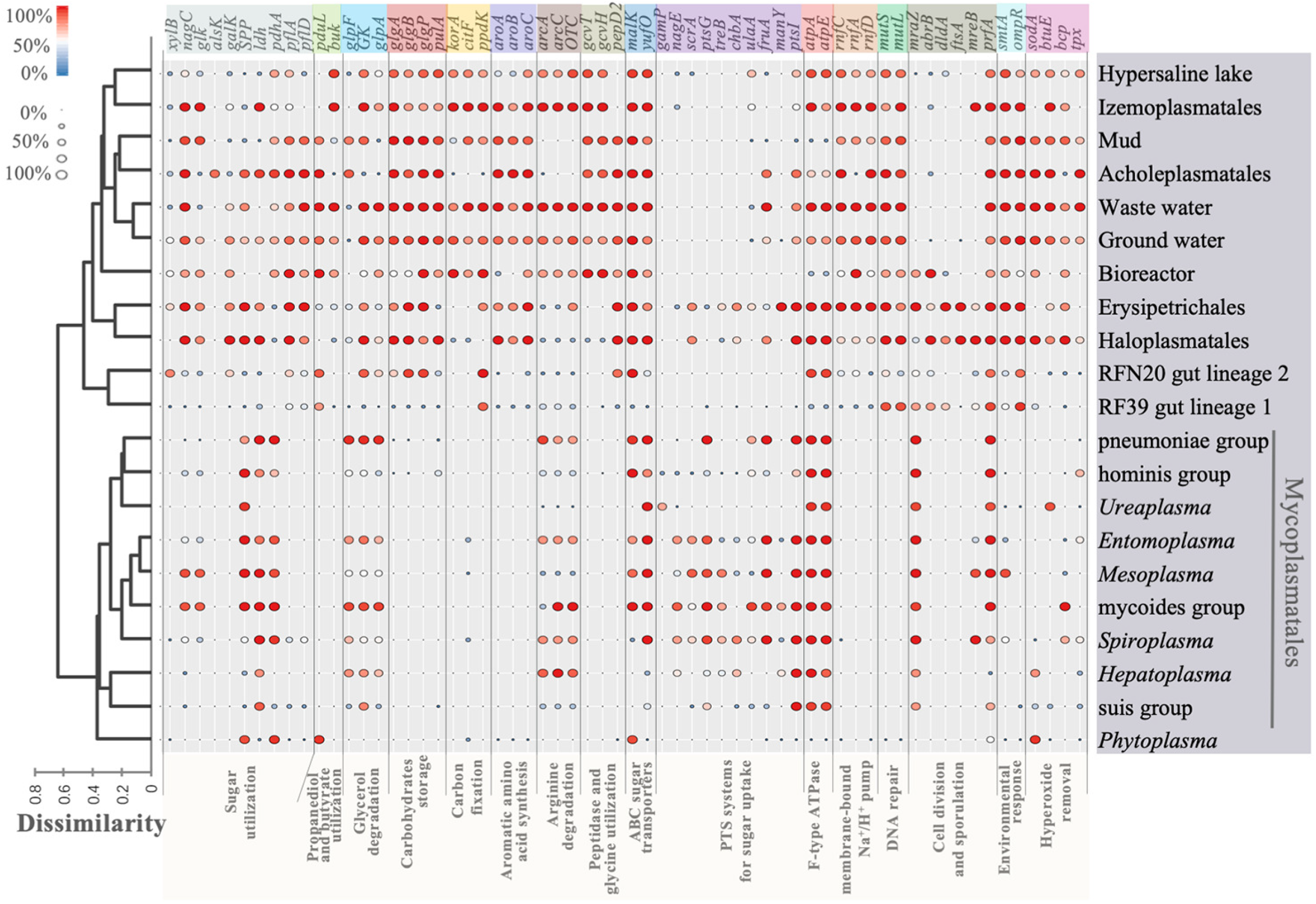
Distribution of genes and pathways in the Tenericutes lineages. Tenericutes lineages were grouped using an agglomerative hierarchical clustering on the basis of the distribution of COGs within each group. The color and size of each dot represent the percentage of genomes within each lineage that carries the gene. The functions of these genes are shown in Table S1.

Aromatic biosynthesis pathway was lost in Mycoplasmatales, indicating their complete dependence on hosts for aromatic amino acids. Acquisition of amino acids by some environmental Tenericutes was likely conducted by peptidases (*pepD2*) and cross-membrane oligopeptide transporters. Glycine was also probably an important carbon and nitrogen source for the environmental Tenericutes, as a high percentage of their genomes (76.3%-100%) contained the glycine cleavage genes *gcvT* and *gcvH*.

Glycerol is a key intermediate between sugar and lipid metabolisms and is imported by a facilitation factor GlpF. Phosphorylation of glycerol by a glycerol kinase (GK) is followed by oxidation to dihydroxyacetone phosphate (DHAP) by glycerol-3-phosphate (G3P) dehydrogenase (GlpD), which is further metabolized in the glycolysis pathway (26). More than 95% of the genomes of *Mesoplasma*, pneumoniae, mycoides and wastewater groups contained the *glpD* gene; in contrast, *Phytoplasma* and *Ureaplasma* genomes lacked a *glpD* gene. 62% of RFN20 genomes harbored the *glpD* gene, while it was only found in 2% of RF39. RF39 genomes also lacked the GK-encoding gene, which suggests that RF39 cannot utilize glycerol from diet or the gut membrane. Hydrogen peroxide (H_2_O_2_) is a by-product of G3P oxidation, and has deleterious effects on epithelial surfaces in humans and animals (27). On the other hand, these H_2_O_2_ catabolism genes were more frequently identified in uncultured environmental Tenericutes (Fig. 3).

The DNA mismatch repair machinery components MutS and MutL were almost entirely absent from Mycoplasmatales and *Phytoplasma* genomes. RFN20 genomes also had a low percentage of the DNA repairing genes (33.3% for *mutS* and 57.1% for *mutL)*. This lack of DNA repairing genes might have generated more mutants in small asexual microbial populations capable of adapting to new environments due to Muller’s ratchet effect (28).

In *Mycoplasma* species as in mitochondria, tRNA anticodon base U34 can pair with any of the four bases in codon family boxes (29). To makes this ability more efficient U34 is modified in some organisms by enzymes using a carboxylated S-adenosylmethionine. The SmtA enzyme, also known as CmoM, is a methyltransferase that adds a further methyl group to U34 modified tRNA for precise decoding of mRNA and rapid growth (30, 31). The high frequency of *smtA* gene in the environmental Tenericutes genomes indicates a capacity to regulate their growth under various conditions. OmpR is a two-component regulator tightly associated with a histidine kinase/phosphatase EnvZ for regulatory response to environmental osmolarity changes(32). Its presence in most of the environmental Tenericutes genomes (>70.4%) suggests its involvement in regulating stress responses in these organisms. The genomes of two gut lineages RFN20 and RF39 also contained a high percentage of the *ompR* gene. In contrast, almost all Mycoplasmatales and *Phytoplasma* genomes lacked the *ompR* gene.

The cell division/cell wall cluster transcriptional repressor MraZ can negatively regulate cell division of Tenericutes (33). The *mraZ* gene that is thus responsible for dormancy of bacteria is conserved in *Erysipelotrichales* and *Mycoplasmatales*. Further studies are needed to examine whether this gene can be targeted to control pathogenicity of the bacteria in the two orders.

The Rnf proton pump system evolved in anoxic condition and is employed by anaerobes to generate proton gradients for energy conservation (34). In single-membrane Tenericutes, proton gradients can hardly be established by the Rnf system due to the leakage of protons directly to the environment. However, this system was well preserved in genomes from Izemoplasmatales and the wastewater group. The Rnf system in these species was likely used for pumping protons out of the cell to balance cytoplasmic pH.

### Metabolic model of gut lineages RFN20 and RF39

A recent study reported the genome features of RFN20 and RF39, the two main clades comprising uncultured Tenericutes (16). The major findings on these two lineages were their small genomes and the lack of several amino acid biosynthesis pathways. After correction for genome completeness in this study, we found that the RF39 genomes were indeed significantly smaller than those of RFN20 genomes (t-test; *p=*0.0012). We selected four nearly complete genomes of RFN20 and RF39 for annotation and elaborated their metabolic potentials (Table 1). The genome sizes were between 1.5 Mb-1.9 Mb, smaller than those from *Sharpea azabuensis* belonging to the order Erysipelotrichales. We built a schematic metabolic map for the representative RFN20 and RF39 species on the basis of the KEGG and COG annotation results. The two lineages were predicted to be acetogens since the four genomes encoded genes for acetate production (Fig. 4). We hypothesize that sugars are imported from the environment by ABC sugar transporters, while autotrophic CO_2_ fixation might occur via carboxylation of acetyl-CoA to pyruvate by the pyruvate:ferredoxin oxidoreductase (PFOR). Glycerol is imported and enters glycerophospholipid metabolism, which results in cardiolipin biosynthesis instead of fermentation through the EMP pathway. In some pathogenic mycoplasmas, glycerol can be taken into central carbon metabolism (26), as mentioned above.

**Table 1.**
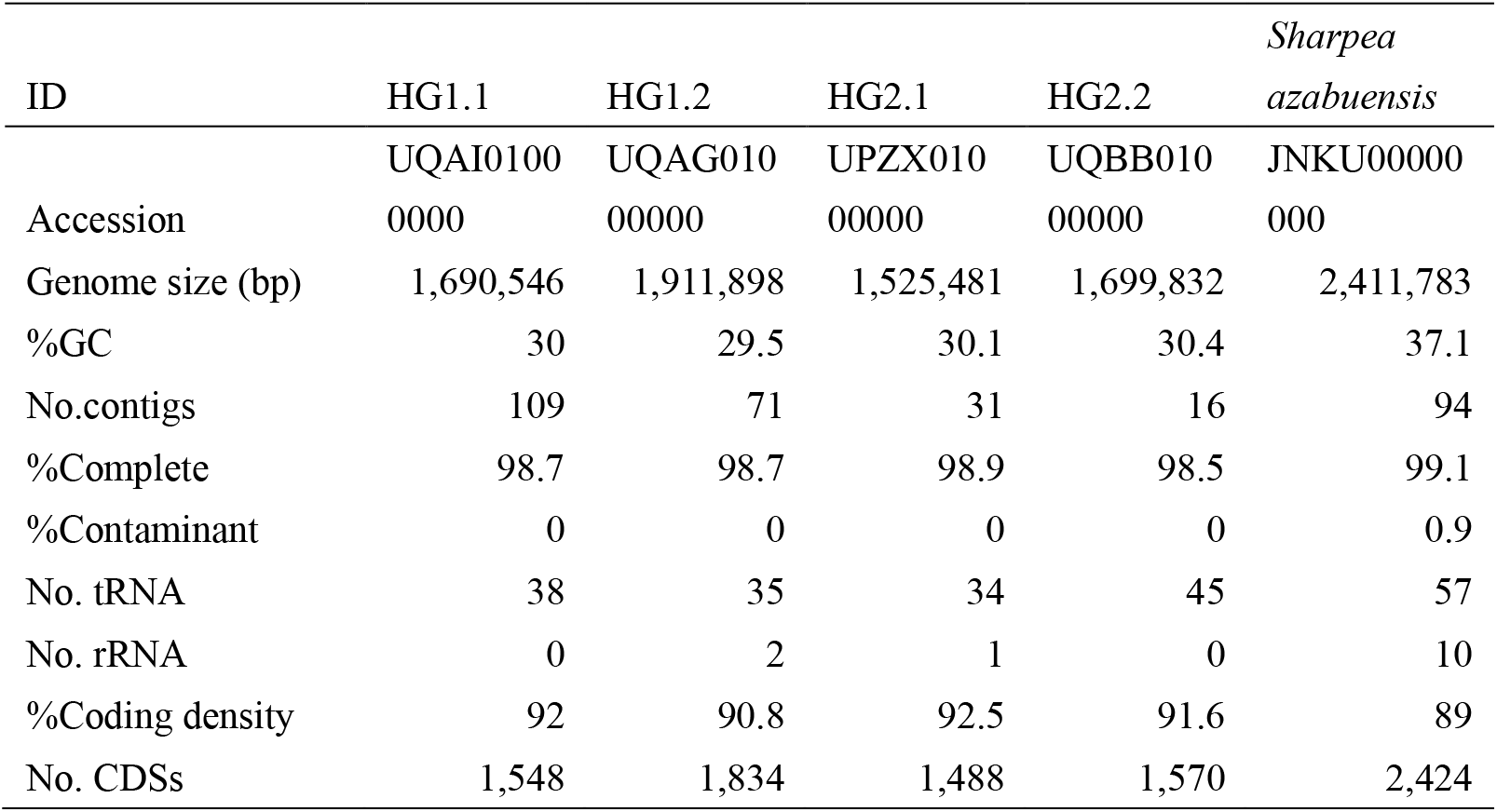
Representative genomes of RFN20 and RF39. RF39 (HG1) was represented by HG1.1 and HG1.2 from the Tenericutes downloaded from NCBI; RFN20 (HG2) was represented by HG2.1 and HG2.2. *S. azabuensis* was a species in *Erysipetrichales*.

**Figure 4.**
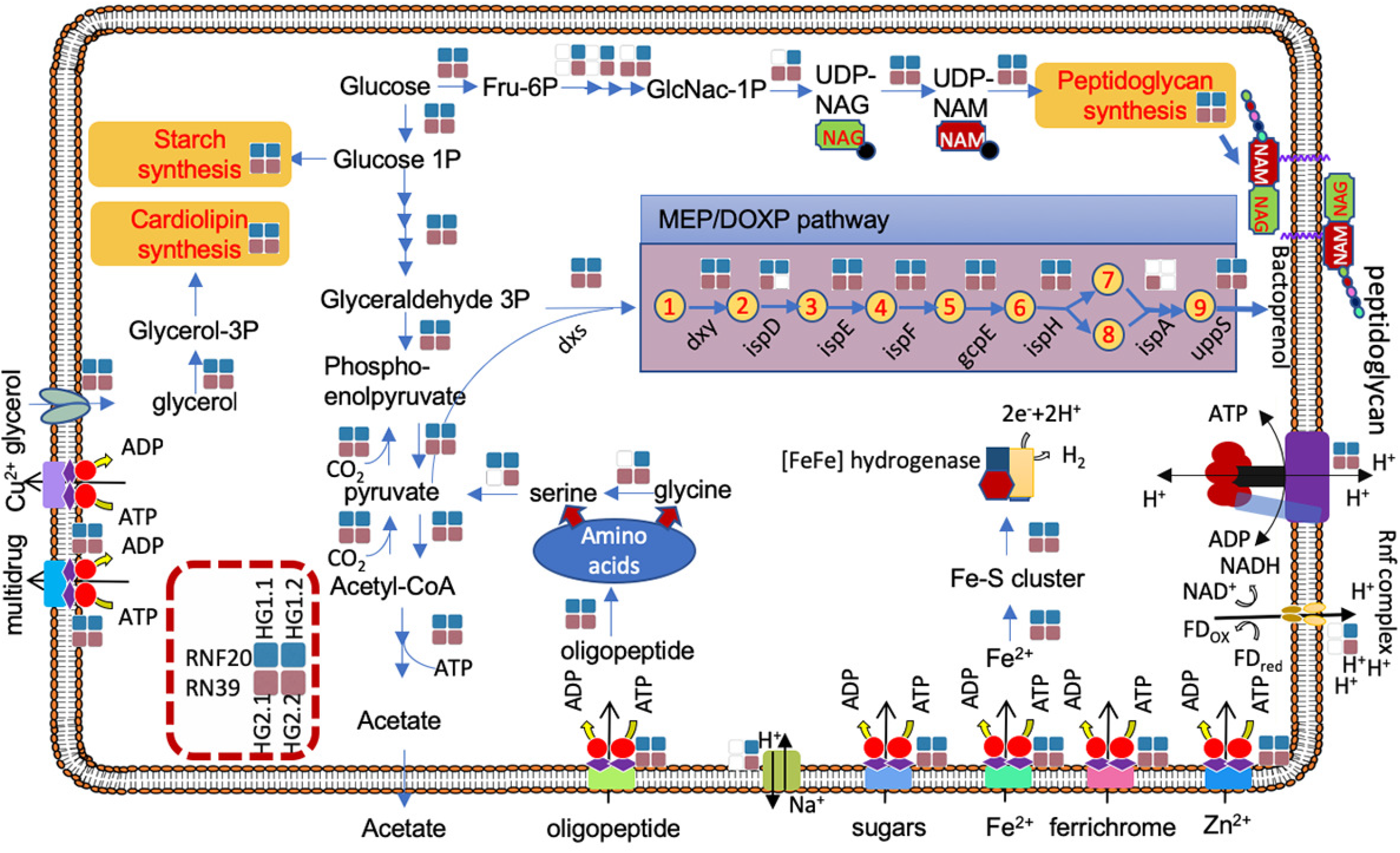
Schematic metabolism of RFN20 and RF39 Metabolic models predicted by using gene annotation results of four representative genomes of RFN20 and RF39 (see Table 1). Solid squares indicate presence of the genes responsible for a step or a pathway. The products depicted in the MEP/DOXP pathway are 1-deoxy-xylulose 5-P, 2-C-methyl-D-erythritol 4-P, 4-(Cytidine 5’-PP)-2-C-methyl-erythritol, 2-P-4-(cytidine 5’-PP)-2-C-methyl-erythritol, 2-C-methyl-erythritol 2,4-PP, 1-hydroxy-2-methyl-2-butenyl 4-PP, dimethylallyl-PP, isopentenyl-PP, and farnesyl-PP.

RFN20 and RF39 are probably mixotrophic since CO_2_ can be fixed to pyruvate and stored as starch, while central carbon metabolism is also connected with amino acid metabolism. After uptake of oligopeptides by the App ABC transporter system, an endo-oligopeptidase encoded by *pepF* yields amino acids for protein synthesis. Glycine and serine might feed into pyruvate metabolism. The peptidoglycan biosynthesis pathway was found to be complete in all four RFN20 and RF39 genomes here considered, but two genomes, namely HG1.1 and HG2.1 (Table 1), lacked the genes encoding the enzymes for UDP-N-acetylglucosamine (UDP-NAG) synthesis. Instead, these genomes harbored all the genes required for the subsequent synthesis steps to generate extracellular peptidoglycan. *murG* and *mraY* genes, which are involved in integration of UDP-NAG and UDP-N-acetylmuramate (UDP-NAM) into the peptidoglycan unit, respectively, were identified in the four genomes. With the addition of an oligopeptide, the peptidoglycan unit is secreted into the cell surface with the assistance of bactoprenol (C55 isoprenoid alcohol) (35, 36), which is formed by condensation of eight isopentenyl-diphosphate (IPP) units and one farnesyl-diphosphate (FPP). The *uppS* gene responsible for the bactoprenol formation was identified in the four RFN20 and RF39 genomes (37). In bacteria, IPP can be synthesized by several metabolic steps. All the genomes contained the genes encoding the respective enzymes involved in the intermediate steps of IPP and dimethylallyl diphosphate (DPP) synthesis through MEP/DOXP pathway, except for *ispD* gene in one genome (Fig. 4). However, the polyprenyl synthetase gene (*ispA*), which is essential in the formation of FPP, was missing in three of the genomes. Given the loss of the *ispA* gene, the source of FPP for bactoprenol synthesis is unclear. Overall, 86.9% and 14.3% of the RF39 and RFN20 genomes contained the *mraY* gene, respectively, while 68.7% and 5.2% of the RF39 and RFN20 genomes had the *murG* gene, respectively. Therefore, most of the RFN20 genomes collected in this study lacked the complete pathway for peptidoglycan synthesis. The two essential genes for peptidoglycan synthesis were only frequently detected in Tenericutes genomes from the bioreactor group (75.0% for both genes) and Erysipelotrichales genomes (80.0% and 60.0% for *mraY* and *murG*, respectively). Therefore, the capacity of peptidoglycan synthesis is possibly deteriorating in the gut lineages, as a potential adaptive strategy to the gut environment. Similarly, the *H. contractile* was reported to possess the peptidoglycan synthesis genes in its genome (4), although it also lacks a cell wall. Our further examination of the genome found that the *murEF* genes involved in extending the oligopeptide attached on UDP-NAM were absent. Hence, the synthesis of aminosugars NAG and NAM probably served as a mechanism of carbon and nitrogen storage for *H. contractile*.

RFN20 and RF39 are probably hydrogen producers, as the four genomes of HG1 and HG2 had [FeFe]-hydrogenase encoding genes. All the genomes carried the *feo* and *fhu* genes for ferrous iron uptake. Ferrous irons are taken by ABC transporters Feo into the cells when ferrous iron concentration is high in the environment. The Fhu receptor for ferrichrome absorption is required in iron-limiting condition such as the human gut (38). The oxygen-sensitive [FeFe]-hydrogenases contain 4Fe-4S cluster and an H-cluster consisting of several conserved catalytic motifs involved in hydrogen production. Three distinct binding motifs of H-cluster in [FeFe]-hydrogenases, TSCXP, PCX_2_KX_2_E and EXMXCXGGCX_2_C (39), were present in the five hydrogenases encoded by all the four genomes (Fig. S2). However, three of the hydrogenases from HG1 and HG2 harbor specific sites that differ from the others in some of the active sites. We have identified several orthologs with these distinct amino acids in the conserved motifs. These [FeFe]-hydrogenases formed a novel cluster in the phylogenetic tree. HG2.1 genome harbored two copies of the [FeFe]-hydrogenase genes, which were diversified as shown by their positions in the phylogenetic tree and the differences in conserved catalytic sites (Fig. S2). In the human gut, three groups of [FeFe]-hydrogenases have been detected, and were proposed to be involved in methanogenesis, acetogenesis and sulfate reduction (40). Lignocellulose-feeding termites also produce a high concentration of hydrogen in their guts, probably for degradation of wood (41). Therefore, the HG1 and HG2 gut lineages are probably important for maintenance of a healthy gut microbial ecosystem and degradation of recalcitrant carbon.

As indicated by the phylogenomics tree, there is a high genomic variation within the RFN20 and RF39 lineages. Therefore, the predicted lifestyle of RFN20 and RF39 may vary among human populations. For example, 68.7% and 76.2% of RF39 and RFN20 genomes, respectively, harbored the *uppS* gene for bactoprenol synthesis. However, the lack of high-quality, isolate genomes representing these lineages hinders the evaluation of their dynamics and evolutionary processes in the human gut.

In this study, the genomic features of RFN20 and RF39 were shown to be highly dynamic among genomes from different sources. RF39 genomes lacked most of the genes for carbohydrate storage but maintained *mutSL* genes involved in DNA repair (Fig. 3). Except for this, there were no major differences between the two lineages, although a previous study claimed that RF39 were prone to be autotrophic (16). In deep-sea isopod gut, we also identified two types of Tenericutes bacteria, *Mycoplasma* sp. Bg1 and Bg2 (6). M. sp. Bg1 was able to degrade sialic acids probably by attachment to the host gut surface. The co-existence of two Tenericutes lineages in human and animal intestinal tracts is still enigmatic and warrants further investigations using microscopy and transcriptomics methods.

In conclusion, our study revealed phylogenetic diversity of the Tenericutes groups and their phylogenomic relationships with Bacilli. In the environmental groups of Tenericutes, we uncovered novel lineages in human guts and marine environments, indicating the lack of environmental representatives for studies on their adaptive strategies and pathogenicity. Our finding of the gut lineages and their metabolic characteristics casts lights into unknown diversified mutualistic Tenericutes in gut microbiome.

## MATERIAL AND METHODS

### Genome collection and quality check

A total of 857 Tenericutes genomes were downloaded from the NCBI database. Three genomes of deep-sea symbiotic Tenericutes were collected from the previous studies (6, 7). Completeness and contamination of the genomes were evaluated by CheckM (v1.0.5) (42). Those with >10% contaminants and <50% complete were removed. To explore variations of GC content in these genomes, GC content within 1-kb genome intervals were calculated. 16S rRNA genes were identified from these genomes using rRNA_HMM with default settings (43), and only those longer than 300 bp were extracted. If there was more than one 16S rRNA gene in a genome, the longest one was selected. The sequences were grouped with an identity cutoff of 99% using CD-HIT (44) and only the longest was retained as the representative. From each order of Bacilli, five genomes (see supplementary file 1) were obtained from the Genome Tree Database (GTDB) (15). They were selected from different families if possible.

### Genome annotation and comparison

The protein coding sequences in the genomes were predicted by Prodigal (v2.6.2) (45) (proteins from Tenericutes in particular were predicted with parameter –g 4). The proteins were then searched against the eggNOG database by eggNOG-mapper (v2) (46) (with parameters --seed_orthorlog_evalue 1e-10), KEGG (24) and COGs (47) databases by Blastp with E-value cutoff of 1e-05 and similarity threshold of 40%. The functions of essential COGs belonging to Tenericutes were referred to those for a synthetic bacterium JCVI-Syn3.0 with a minimal genome (48).

The collected Tenericutes genomes were grouped by taxonomy and source (supplementary file 1). The percentage of the KEGG genes and COGs in the genomes of each group was calculated. This was also accomplished for *Erysipelotrichales* genomes. To filter low-frequency genes, at least one of the groups had a target gene in >30% of the genomes. The percentages of the genes used for Bray-Curtis dissimilarity estimates were calculated using the COG frequency table. AHC analysis was conducted using the pairwise dissimilarities between groups. A Mann-Whitney test was performed using the percentages of COGs and KEGG genes between the AHC clusters. The KEGG genes with *p* value <0.01 were clustered into functional modules on the KEGG website (www.kegg.jp).

### Phylogenetic and phylogenomic analyses

The analyses on the datasets of 16S rRNA gene amplicons from marine samples were described in our previous study (49). The representative reads of Tenericutes OTUs were recruited for this study. Raw metagenomic data from Tara Ocean project were checked by FastQC (version 0.11.4). Reads with low quality bases (PHRED quality score < 20 over 70% of the reads) were removed using the NGS QC Toolkit (50). The quality-filtered reads were merged using PEAR (v0.9.5) (51) and those 16S rRNA fragments >140 bp were identified and extracted with rRNA_HMM (43). After taxonomic classification of the fragments using the Ribosomal Database Project (RDP) classifier version 2.2 against the SILVA 128 database (52, 53), those belonging to Tenericutes were collected for the following phylogenetic study.

The 16S rRNA genes from the genomes, the amplicons and the Tara project were first clustered by MUSCLE (v3.8) (54) and then trimmed by trimAl v1.4 (automated1) (55). The ML phylogenetic tree of 16S rRNA genes was built by IQ-TREE (v1.6.10) (56, 57) (with parameters -m GTR+F+R10 -alrt 1000 -bb 1000). Conserved proteins of the Tenericutes genomes were identified by AMPHORA2 (58). A total of 31 conserved proteins were used to construct the phylogenomic tree for Tenericutes. The conserved proteins were aligned with MUSCLE (v3.8)(54), concatenated and then trimmed with trimAl (v1.4) (automated1) (55). The conserved proteins from *Syntrophomonas wolfei* (NC_008346), *Thermacetogenium phaeum* (NC_018870) and *Desulfallas geothermicus* (NZ_FOYM01000001) were combined with the dataset of Tenericutes as an outgroup. The phylogenomics tree for Tenericutes was built by IQ-TREE (v1.6.10) (56, 57) (with parameters -m LG+F+R10 -alrt 1000 -bb 1000). The phylogenomic tree for Bacilli and Tenericutes was constructed first with IQ-TREE (v1.6.10) using the same settings as that for the phylogenomics tree of Tenericutes and then with RAxML 8.1.21 using PROTGAMMA+BLOSUM62 model with 100 bootstrap replicates.

### Prediction of metabolic models of RFN20 and RF39

Four genomes were selected from the downloaded genomes of Tenericutes to represent RFN20 and RF39 with respect to their high genome completeness. The protein-coding sequences were predicted by Prodigal (v2.6.2) (45) as mentioned above. The proteins were then searched against COG database (47) by Blastp (59) with an E-value cutoff of 1e-05. KEGG annotation was conducted using the online BlastKOALA tool (24).

## ACKNOWLEDGEMENTS

This study was supported by the National Key Research and Development Program of China (2016YFC0302504 and 2018YFC0310005). AA and RDF are supported by European Molecular Biology Laboratory core funds. We thank Shriya Raj for comments and feedback on the manuscript.

Y.W., A.D. and L.S.H. designed the study; Y.W., J.M.H., and Y.L.Z. performed the bulk of the phylogenomic analyses; A.A. and R.D.F. contributed data for analysis;

Y.W. wrote the manuscript. All of us contributed to manuscript revisions.

The authors declare that there is no conflict of interest.

## REFERENCES

1. Razin S, Herrmann R. 2002. Molecular biology and pathogenicity of mycoplasmas. Springer, Boston, MA.

2. Antunes A, Rainey FA, Wanner G, Taborda M, Pätzold J, Nobre MF, da Costa MS, Huber R. 2008. A new lineage of halophilic, wall-less, contractile bacteria from a brine-filled deep of the Red Sea. J Bacteriol 190:3580–3587.

3. Skennerton CT, Haroon MF, Briegel A, Shi J, Jensen GJ, Tyson GW, Orphan VJ. 2016. Phylogenomic analysis of Candidatus ‘Izimaplasma’ species: free-living representatives from a Tenericutes clade found in methane seeps. ISME J 10:2679–2692.

4. Antunes A, Alam I, El Dorry H, Siam R, Robertson A, Bajic VB, Stingl U. 2011. Genome sequence of Haloplasma contractile, an unusual contractile bacterium from a deep-sea anoxic brine lake. J Bacteriol 193:4551–4552.

5. Wang YJ, Stingl U, Anton-Erxleben F, Geisler S, Brune A, Zimmer M. 2004. “*Candidatus* Hepatoplasma crinochetorum,” a new, stalk-forming lineage of Mollicutes colonizing the midgut glands of a terrestrial isopod. Appl Environ Microb 70:6166–6172.

6. Wang Y, Huang JM, Wang SL, Gao ZM, Zhang AQ, Danchin A, He LS. 2016. Genomic characterization of symbiotic mycoplasmas from the stomach of deep-sea isopod bathynomus sp. Environ Microbiol 18:2646–2659.

7. He L-S, Zhang P-W, Huang J-M, Zhu F-C, Danchin A, Wang Y. 2018. The enigmatic genome of an obligate ancient Spiroplasma symbiont in a hadal holothurian. Appl Environ Microbiol 84:e01965–01917.

8. Sullam KE, Essinger SD, Lozupone CA, O’Connor MP, Rosen GL, Knight R, Kilham SS, Russell JA. 2012. Environmental and ecological factors that shape the gut bacterial communities of fish: a meta-analysis. Mol Ecol 21:3363–3378.

9. Yun JH, Roh SW, Whon TW, Jung MJ, Kim MS, Park DS, Yoon C, Nam YD, Kim YJ, Choi JH, Kim JY, Shin NR, Kim SH, Lee WJ, Bae JW. 2014. Insect gut bacterial diversity determined by environmental habitat, diet, developmental stage, and phylogeny of host. Appl Environ Microbiol 80:5254–5264.

10. Aceves AK, Johnson P, Bullard SA, Lafrentz S, Arias CR. 2018. Description and characterization of the digestive gland microbiome in the freshwater mussel Villosa nebulosa (Bivalvia: Unionidae). J Molluscan Studies 84:240–246.

11. Almeida A, Mitchell AL, Boland M, Forster SC, Gloor GB, Tarkowska A, Lawley TD, Finn RD. 2019. A new genomic blueprint of the human gut microbiota. Nature 568:499–504.

12. Moran NA. 2002. Microbial minimalism: Genome reduction in bacterial pathogens. Cell 108:583–586.

13. Lo W-S, Gasparich GE, Kuo C-H. 2018. Convergent evolution among ruminant-pathogenic mycoplasma involved extensive gene content changes. Genome Biol Evol 10:2130–2139.

14. Chernov VM, Chernova OA, Mouzykantov AA, Medvedeva ES, Baranova NB, Malygina TY, Aminov RI, Trushin MV. 2018. Antimicrobial resistance in mollicutes: known and newly emerging mechanisms. FEMS Microbiol Lett 365.

15. Parks DH, Chuvochina M, Waite DW, Rinke C, Skarshewski A, Chaumeil PA, Hugenholtz P. 2018. A standardized bacterial taxonomy based on genome phylogeny substantially revises the tree of life. Nature Biotechnol 36:996–1004.

16. Nayfach S, Shi ZJ, Seshadri R, Pollard KS, Kyrpides NC. 2019. New insights from uncultivated genomes of the global human gut microbiome. Nature 568:505–510.

17. Zhang LT, Huang XF, Xue B, Peng QH, Wang ZS, Yan TH, Wang LZ. 2015. Immunization against rumen methanogenesis by vaccination with a new recombinant protein. PLoS ONE 10:e0140086.

18. Cheng X-Y, Wang Y, Li J-Y, Yan G-Y, He L-S. 2019. Comparative analysis of the gut microbial communities between two dominant amphipods from the Challenger Deep, Mariana Trench. Deep Sea Res I 151:103081.

19. Sunagawa S, Coelho LP, Chaffron S, Kultima JR, Labadie K, Salazar G, Djahanschiri B, Zeller G, Mende DR, Alberti A, Cornejo-Castillo FM, Costea PI, Cruaud C, d’Ovidio F, Engelen S, Ferrera I, Gasol JM, Guidi L, Hildebrand F, Kokoszka F, Lepoivre C, Lima-Mendez G, Poulain J, Poulos BT, Royo-Llonch M, Sarmento H, Vieira-Silva S, Dimier C, Picheral M, Searson S, Kandels-Lewis S, Bowler C, de Vargas C, Gorsky G, Grimsley N, Hingamp P, Iudicone D, Jaillon O, Not F, Ogata H, Pesant S, Speich S, Stemmann L, Sullivan MB, Weissenbach J, Wincker P, Karsenti E, Raes J, Acinas SG, Bork P. 2015. Structure and function of the global ocean microbiome. Science 348:1261359.

20. Pitta DW, Parmar N, Patel AK, Indugu N, Kumar S, Prajapathi KB, Patel AB, Reddy B, Joshi C. 2014. Bacterial diversity dynamics associated with different diets and different primer pairs in the rumen of kankrej cattle. PLoS ONE 9:e111710.

21. Shimoji Y, Yokomizo Y, Sekizaki T, Mori Y, Kubo M. 1994. Presence of a capsule in Erysipelothrix-Rhusiopathiae and its relationship to virulence for mice. Infect Imm 62:2806–2810.

22. Soppa J. 2006. From genomes to function: haloarchaea as model organisms. Microbiology 152:585–590.

23. Lyubchenko YL, Shlyakhtenko LS. 1997. Visualization of supercoiled DNA with atomic force microscopy in situ. Proc Natl Acad Sci U S A 94:496–501.

24. Kanehisa M, Goto S. 2000. KEGG: kyoto encyclopedia of genes and genomes. Nucl Acids Res 28:27–30.

25. Tjaden B, Plagens A, Dorr C, Siebers B, Hensel R. 2006. Phosphoenolpyruvate synthetase and pyruvate, phosphate dikinase of *Thermoproteus tenax*: key pieces in the puzzle of archaeal carbohydrate metabolism. Mol Microbiol 60:287–298.

26. Yeh JI, Chinte U, Du S. 2008. Structure of glycerol-3-phosphate dehydrogenase, an essential monotopic membrane enzyme involved in respiration and metabolism. Proc Natl Acad Sci U S A 105:3280–3285.

27. Blotz C, Stulke J. 2017. Glycerol metabolism and its implication in virulence in *Mycoplasma*. FEMS Microbiol Rev 41:640–652.

28. Andersson DI, Hughes D. 1996. Muller’s ratchet decreases fitness of a DNA-based microbe. Proc Natl Acad Sci U S A 93:906–907.

29. Grosjean H, Westhof E. 2016. An integrated, structure- and energy-based view of the genetic code. Nucl Acids Res 44:8020–8040.

30. Sakai Y, Miyauchi K, Kimura S, Suzuki T. 2016. Biogenesis and growth phase-dependent alteration of 5-methoxycarbonylmethoxyuridine in tRNA anticodons. Nucl Acids Res 44:509–523.

31. Yamanaka K, Ogura T, Niki H, Hiraga S. 1995. Characterization of the smtA gene encoding an S-adenosylmethionine-dependent methyltransferase of Escherichia coli. FEMS Microbiol Lett 133:59–63.

32. Cai SJ, Inouye M. 2002. EnvZ-OmpR interaction and osmoregulation in Escherichia coli. J Biol Chem 277:24155–24161.

33. Eraso JM, Markillie LM, Mitchell HD, Taylor RC, Orr G, Margolin W. 2014. The highly conserved MraZ protein is a transcriptional regulator in Escherichia coli. J Bacteriol 196:2053–2066.

34. Schuchmann K, Muller V. 2014. Autotrophy at the thermodynamic limit of life: a model for energy conservation in acetogenic bacteria. Nat Rev Microbiol 12:809–821.

35. Thorne KJ, Kodicek E. 1966. The structure of bactoprenol, a lipid formed by lactobacilli from mevalonic acid. Biochem J 99:123–127.

36. Manat G, Roure S, Auger R, Bouhss A, Barreteau H, Mengin-Lecreulx D, Touzé T. 2014. Deciphering the metabolism of undecaprenyl-phosphate: the bacterial cell-wall unit carrier at the membrane frontier. Microb Drug Ris 20:199–214.

37. Mostafavi AZ, Lujan DK, Erickson KM, Martinez CD, Troutman JM. 2013. Fluorescent probes for investigation of isoprenoid configuration and size discrimination by bactoprenol-utilizing enzymes. Bioorganic Med Chem 21:5428–5435.

38. Wooldridge KG, Williams PH. 1993. Iron uptake mechanisms of pathogenic bacteria. FEMS Microbiol Rev 12:325–348.

39. Mulder David W, Shepard Eric M, Meuser Jonathan E, Joshi N, King Paul W, Posewitz Matthew C, Broderick Joan B, Peters John W. 2011. Insights into [FeFe]-hydrogenase structure, mechanism, and maturation. Structure 19:1038–1052.

40. Wolf PG, Biswas A, Morales SE, Greening C, Gaskins HR. 2016. H-2 metabolism is widespread and diverse among human colonic microbes. Gut Microbes 7:235–245.

41. Ballor NR, Leadbetter JR. 2012. Patterns of [FeFe] hydrogenase diversity in the gut microbial communities of lignocellulose-feeding higher termites. Appl Environ Microb 78:5368–5374.

42. Parks DH, Imelfort M, Skennerton CT, Hugenholtz P, Tyson GW. 2015. CheckM: assessing the quality of microbial genomes recovered from isolates, single cells, and metagenomes. Genome Res 25:1043–1055.

43. Huang Y, Gilna P, Li W. 2009. Identification of ribosomal RNA genes in metagenomic fragments. Bioinformatics 25:1338–1340.

44. Fu LM, Niu BF, Zhu ZW, Wu ST, Li WZ. 2012. CD-HIT: accelerated for clustering the next-generation sequencing data. Bioinformatics 28:3150–3152.

45. Hyatt D, Locascio PF, Hauser LJ, Uberbacher EC. 2012. Gene and translation initiation site prediction in metagenomic sequences. Bioinformatics 28:2223–2230.

46. Huerta-Cepas J, Forslund K, Coelho LP, Szklarczyk D, Jensen LJ, Von MC, Bork P. 2016. Fast genome-wide functional annotation through orthology assignment by eggNOG-mapper. Mol Biol Evol 34:2115.

47. Galperin MY, Makarova KS, Wolf YI, Koonin EV. 2015. Expanded microbial genome coverage and improved protein family annotation in the COG database. Nucl Acids Res 43:261–269.

48. Hutchison CA, 3rd, Chuang RY, Noskov VN, Assad-Garcia N, Deerinck TJ, Ellisman MH, Gill J, Kannan K, Karas BJ, Ma L, Pelletier JF, Qi ZQ, Richter RA, Strychalski EA, Sun L, Suzuki Y, Tsvetanova B, Wise KS, Smith HO, Glass JI, Merryman C, Gibson DG, Venter JC. 2016. Design and synthesis of a minimal bacterial genome. Science 351:aad6253.

49. Li W-L, Huang J-M, Zhang P-W, Cui G-J, Wei Z-F, Wu Y-Z, Gao Z-M, Han Z, Wang Y. 2019. Periodic and spatial spreading of alkanes and Alcanivorax bacteria in deep waters of the Mariana Trench. Appl Environ Microbiol 85:e02089–02018.

50. Patel RK, Jain M. 2012. NGS QC toolkit: A toolkit for quality control of next generation sequencing data. PLoS ONE 7:e30619.

51. Zhang J, Kobert K, Flouri T, Stamatakis A. 2014. PEAR: a fast and accurate Illumina Paired-End reAd mergeR. Bioinformatics 30:614.

52. Caporaso JG, Bittinger K, Bushman FD, Desantis TZ, Andersen GL, Knight R. 2010. PyNAST: a flexible tool for aligning sequences to a template alignment. Bioinformatics 26:266–267.

53. Wang Q, Garrity GM, Tiedje JM, Cole JR. 2007. Naïve Bayesian Classifier for Rapid Assignment of rRNA Sequences into the New Bacterial Taxonomy. Appl Environ Microbiol 73:5261.

54. Edgar RC. 2004. MUSCLE: multiple sequence alignment with high accuracy and high throughput. Nucl Acids Res 32:1792–1797.

55. Capellagutiérrez S, Sillamartínez JM, Gabaldón T. 2009. trimAl: a tool for automated alignment trimming in large-scale phylogenetic analyses. Bioinformatics 25:1972–1973.

56. Lam-Tung N, Schmidt HA, Arndt VH, Bui Quang M. 2015. IQ-TREE: a fast and effective stochastic algorithm for estimating maximum-likelihood phylogenies. Mol Biol Evol 32:268–274.

57. Kalyaanamoorthy S, Minh BQ, Wong TKF, Haeseler AV, Jermiin LS. 2017. ModelFinder: fast model selection for accurate phylogenetic estimates. Nature Meth 14: 587–589.

58. Wu M, Scott AJ. 2012. Phylogenomic analysis of bacterial and archaeal sequences with AMPHORA2. Bioinformatics 28:1033–1034.

59. Altschul SF, Gish W, Miller W, Myers EW, Lipman DJ. 1990. Basic local alignment search tool. J Mol Biol 215:403–410.

